# The mineralocorticoid receptor forms higher order oligomers upon DNA binding

**DOI:** 10.1101/2023.01.26.525752

**Authors:** Gregory Fettweis, Thomas A. Johnson, Brian Almeida-Prieto, Diego M. Presman, Gordon L. Hager, Diego Alvarez de la Rosa

**Author notes:** corresponding authors: G.L.H.,; D.A.d.l.R. Present address: Laboratory of Gene Expression and Cancer, GIGA-Molecular Biology of Disease, University of Liège, Liège, Belgium. **Summary:** Closely related corticosteroid receptors adopt divergent quaternary structures in their active conformations but still interact to determine aldosterone and glucocorticoid signaling.

## Abstract

The prevailing model of steroid hormone nuclear receptor function assumes ligand-induced homodimer formation followed by binding to DNA hormone response elements (HREs). This model has been challenged by evidence showing that the glucocorticoid receptor (GR) forms tetramers upon ligand and DNA binding, which then drive receptor-mediated gene transactivation and transrepression. GR and the closely-related mineralocorticoid receptors (MR) interact to transduce corticosteroid hormone signaling, but whether they share the same quaternary arrangement is unknown. Here, we used a fluorescence imaging technique, Number & Brightness, to study oligomerization in a cell system allowing real-time analysis of receptor-DNA interactions. Agonist-bound MR forms tetramers in the nucleoplasm and higher order oligomers upon binding to HREs. Antagonists form intermediate quaternary arrangements, suggesting that large oligomers are essential for function. Divergence between MR and GR quaternary structure is driven by different functionality of known and new multimerization interfaces, which does not preclude formation of heteromers. Thus, influencing oligomerization may be important to selectively modulate corticosteroid signaling.

## INTRODUCTION

The mineralocorticoid and glucocorticoid receptors (MR and GR, respectively) are members of the steroid hormone receptor subfamily of nuclear receptors (NR3C). Steroid receptors share a common modular protein architecture (Green and Chambon, 1987) that includes a highly-divergent N-terminal domain (NTD), a highly conserved DNA-binding domain and a moderately conserved ligand-binding domain (DBD and LBD) (Grossmann et al., 2022). The NTD of both MR and GR is intrinsically disordered but both contain divergent ligand-independent transcription activation function 1 domains (AF-1), that provide an interaction surface for a diversity of transcriptional co-regulators (Fuse et al., 2000; Grossmann et al., 2022; Lavery and McEwan, 2005; Tallec et al., 2003). The highly conserved DBD between MR and GR imparts essentially indistinguishable DNA binding specificity (Hudson et al., 2014). The LBD similarity between both receptors confers promiscuous ligand activation for MR, binding both mineralocorticoids such as aldosterone or glucocorticoids such as cortisol or corticosterone with similar high affinity (Arriza et al., 1987; Bledsoe et al., 2002; Fagart et al., 2005; Li et al., 2005). MR and GR evolved from a gene duplication event predating the appearance of aldosterone (Baker and Katsu, 2019). MR was then co-opted as a receptor system for a new class of steroid hormones, mineralocorticoids, which regulate mineral and water homeostasis (Rossier et al., 2015). However, MR retained its high-affinity glucocorticoid binding and thus participates in mediating biological responses to both types of hormones (Gomez-Sanchez and Gomez-Sanchez, 2014; Grossmann et al., 2022). Thus, MR and GR have distinct but overlapping physiological functions. Inappropriate activation of MR may mimic or counteract GR actions, promoting obesity and metabolic syndrome (Fallo et al., 2006; Schreier et al., 2022), enhancing inflammation and tissue fibrosis (van der Heijden et al., 2022) or diminishing it (Bigas et al., 2018) in a tissue-specific fashion (Gomez-Sanchez and Gomez-Sanchez, 2014), or modulating brain responses to stress (Paul et al., 2022). MR inhibitors, initially used to treat conditions derived from hyperaldosteronism and situations with excessive water retention have increasingly attracted interest as anti-inflammatory and anti-fibrotic targets (Jaisser and Farman, 2016; Lother et al., 2022), with important applications to treat cardiovascular disease (Bauersachs and Lopez-Andres, 2022) and recently approved indications to treat patients with chronic kidney disease associated with type 2 diabetes ((Barrera-Chimal et al., 2022)). In addition to functional crosstalk between MR and GR, there is considerable evidence pointing towards physical interaction and two-way transcriptional modulation between both receptors (Bigas et al., 2018; Carceller-Zazo et al., 2023; Jimenez-Canino et al., 2016; Liu et al., 1995; Nishi et al., 2004; Pooley et al., 2020; Rivers et al., 2019; Trapp et al., 1994).

Growing interest in the molecular basis of specific MR and GR action and the functional role of their physical interaction make it essential to understand the active conformations of both receptors. The prevailing model of dimers as the final active conformation of steroid receptors has been challenged in the past few years (Jimenez-Panizo et al., 2022; Paakinaho et al., 2019; Presman et al., 2016; Presman and Hager, 2017). Using the Number & Brightness (N&B) assay (Digman et al., 2008) to measure average oligomer size with high spatial resolution in living cells, we have previously reported that agonist-bound GR adopts a dimeric conformation in the nucleoplasm, where the majority of the receptor is soluble, indicating that dimerization precedes high-affinity DNA binding (Presman et al., 2016; Presman et al., 2014). Observation of fluorescently-tagged GR at an array of mouse mammary tumor virus (MMTV) long terminal repeats, containing multiple HRE elements, suggested that the receptor adopts a tetrameric organization upon DNA binding (Presman et al., 2016). The progesterone receptor (PR) adopts a tetrameric conformation regardless of the nuclear compartment, while the androgen receptor (AR) forms larger oligomeric complexes, with an average of 6 subunits in all nuclear compartments (Presman et al., 2016). This paradigm shift in steroid receptor quaternary organization (Fuentes-Prior et al., 2019; Presman and Hager, 2017) and the close evolutionary relationship of GR with MR, in addition to the formation of heterocomplexes between both receptors, bring forward the question of whether MR and GR share a common quaternary structure. In this study, we used the N&B technique to examine MR oligomerization upon ligand binding. We show striking differences between MR and GR, with MR adopting a tetrameric conformation in the nucleoplasm and forming higher order oligomers upon HRE binding. Known or proposed dimerization interfaces conserved between MR and GR have distinct properties in both receptors. In spite of these differences, GR is able to displace MR subunits and incorporate into heterocomplexes. Our results suggest that modulation of quaternary conformation may be an important parameter to take into consideration during development of selective corticosteroid signaling modulators.

## RESULTS

### Agonist-bound MR forms large oligomers at hormone response elements

MR quaternary structure was investigated in a cell derived from murine C127 adenocarcinoma cell line incorporating in their genome a tandem gene array of approximately 200 copies of the mouse mammary tumor virus promoter (MMTV array), which contain hormone response elements (HREs) (McNally et al., 2000). This cell line was further modified using CRISPR/Cas9 to knockout the endogenous expression of GR, as previously described (Paakinaho et al., 2019). Therefore, cells transiently transfected with eGFP-tagged MR allow visualization of the receptor in the nucleoplasm and also enriched at a specific nuclear domain (MMTV array), in the absence of any endogenous GR or MR expression. MR oligomerization was studied using number & brightness (N&B) analysis, a technique that has previously been used to study oligomerization of GR and other steroid receptors in live cells (Presman et al., 2016; Presman et al., 2014). N&B estimates the molecular brightness (ε) of a fluorophore using the first (mean) and second (variance) moments of the intensity fluctuations observed on each pixel of a raster-scan image (Digman et al., 2008). This way, one can obtain the weighted-average brightness (i.e., oligomeric state) of a protein in the entire nucleus or at a specific region such as the MMTV array. As monomeric and dimeric standards, all experiments included a condition with expression of GR truncated after amino acid 525 (GR-N525), which has been shown to exist in monomeric form in the nucleoplasm and to form dimers at the MMTV gene array (Presman et al., 2016). This allowed us to normalize every experiment and calculate MR oligomerization relative to this mutant. Unstimulated cells show partial localization of MR to the nucleus (Fig. 1A), as previously described (Fejes-Toth et al., 1998; Walther et al., 2005). Upon treatment with hormone agonists (either aldosterone or corticosterone), MR fully translocated to the nucleus and produced a bright focus at the MMTV array (Fig. 1A, arrows) (McNally et al., 2000). N&B analysis showed that unstimulated nuclear MR exists as a monomer (ε = 0.96), but reaches an ε of approximately 4 (4.39; Fig. 1B) upon stimulation with a saturating concentration of aldosterone (10 nM), suggesting the formation of a tetramer in the nucleoplasm. N&B analysis of MR expressed from a stably integrated locus also provided an ε near 4, indicating that tetramerization is not an artifact due to transient overexpression (supplementary Fig. S1A). Aldo-stimulated MR concentrated at the MMTV array produces an even higher ε, which approaches 7 (6.92; Fig. 1B), suggesting that higher order oligomerization of MR correlates with chromatin binding and transcriptionally active HREs.

**Figure 1.**
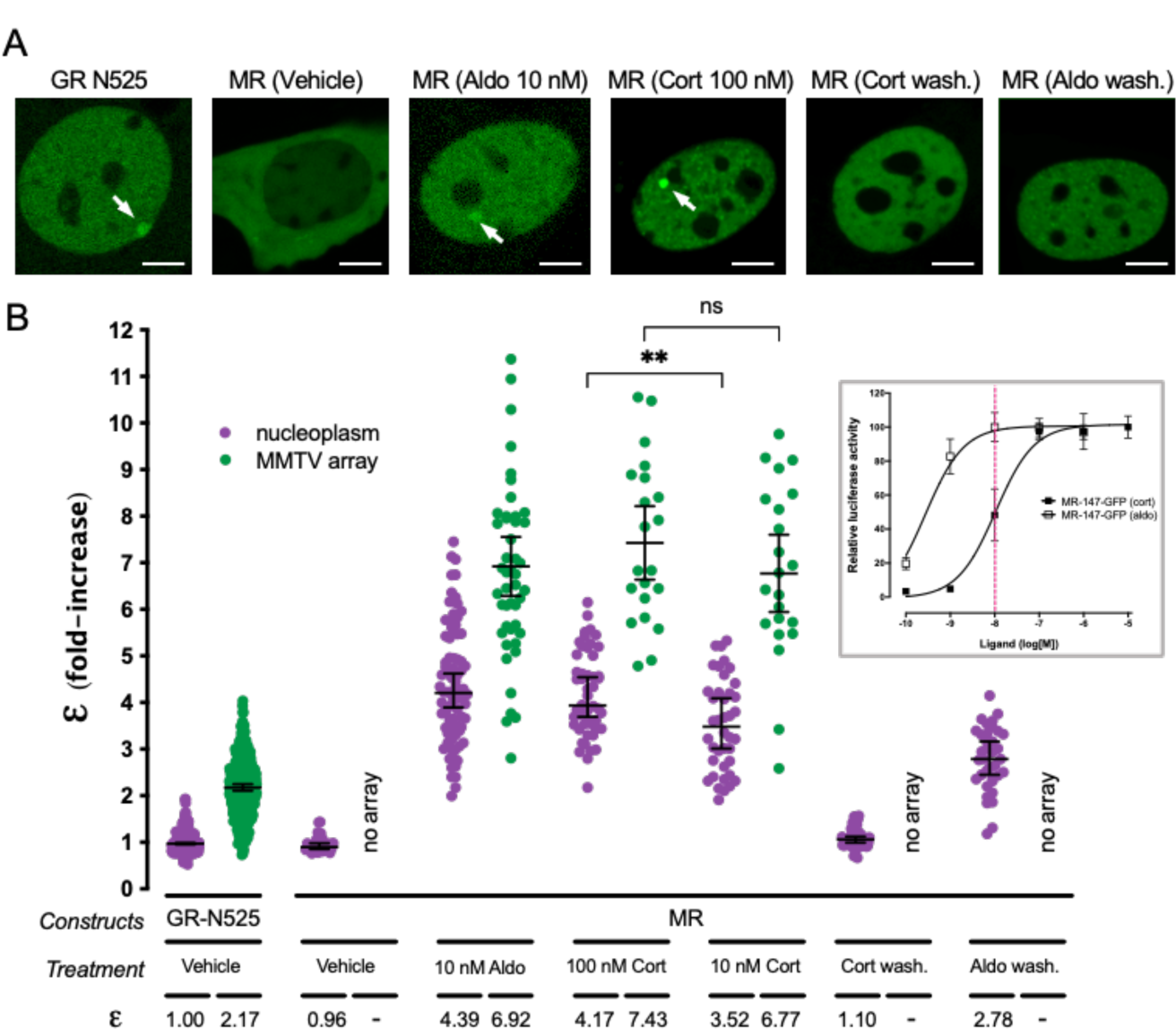
Agonist-bound MR adopts a tetramer conformation in the nucleoplasm and forms higher order oligomers at HREs. (A) Representative images of single cell nuclei expressing GFP-GR-N525 or MR-GFP and treated with vehicle, 10 nM aldosterone (Aldo) or 100 nM corticosterone (Cort). In certain experiments, agonists were washed out after one hour stimulation (wash.). White arrows point to the MMTV array. Scale bars: 5 µm. (B) Normalized molecular brightness (ε). Each point represents a single nucleus (n = 490, 307, 26, 82, 47, 50, 21, 40, 22, 55, 36 cells in each condition, from left to right). Horizontal bars represent mean ± 95% confidence interval (CI). Wash., washout. Statistical analysis for the selected pairs (MR 100 nM CORT vs. 10 nM CORT) was performed using an unpaired t test (**, p < 0.01; n.s., not significant). Inset, MR dose-response gene transactivation curves in response to aldosterone (aldo) and corticosterone (cort). Curves were obtained using wild-type mouse MR transiently transfected in COS7 cells co-expressing a luciferase gene reporter system and treated with the indicated concentrations of hormones for 16h. Dashed red line highlights the difference in MR activity at 10 nM hormone concentration (data adapted from (Jimenez-Canino et al., 2016)).

Since both aldosterone and glucocorticoids are endogenous agonists of MR, we tested whether receptor oligomerization varies as a function of the agonist. To that end, we compared the results obtained with aldosterone to those obtained with saturating concentrations of corticosterone (Cort, 100 nM), a dose that produces full receptor activation (Fig. 1B, inset). Our results show that both agonists produce indistinguishable ε values that are consistent with a tetrameric organization in the nucleoplasm and a higher order oligomer on the MMTV array (Fig. 1A and 1B). To further test the relationship between agonist binding and oligomerization, we took advantage of an important difference between aldosterone and corticosterone. Both hormones bind MR with equal high affinity (*K_d_* ∼ 0.5 nM), but aldosterone is more effective in activating the receptor (EC_50_[aldo] = 0.5 nM vs. EC_50_[cort] = 10 nM). This difference in potency has been ascribed to a higher off-rate of glucocorticoids in the receptor (Lombes et al., 1994).

Therefore, we tested a lower concentration of corticosterone (10 nM), which fully saturates the receptor but produces 50% of the maximum activity (Fig. 1B, inset, vertical dashed line). Under those conditions, nucleoplasmic MR showed a statistically significant lower ε value (ε = 3.52; Fig. 1B), suggesting a correlation between hormone off-rate and oligomerization. In contrast, MR at the MMTV array still showed a high value (ε = 6.77), which is consistent with required ligand binding for high-affinity interaction with HREs (Groeneweg et al., 2014).

We next studied the reversibility of agonist-induced oligomerization of MR. To that end cells were incubated with agonists for 1 hour, followed by hormone washout and an additional 4-hour incubation before recording. Under those conditions, MR still showed full nuclear localization (i.e., negligible nuclear export), but no binding to the MMTV array (Fig. 1A). N&B analysis revealed that after corticosterone washout MR reverted to a monomeric organization in the nucleoplasm (Fig. 1B). In contrast, aldosterone washout produced incomplete reversal, with a nucleoplasmic MR ε value of 2.78 (Fig. 1B), suggesting a mixed population of tetramers with dimers and/or monomers. This could be explained by the shorter off-rate of aldosterone binding to the receptor (Lombes et al., 1994). These results demonstrate that higher order oligomerization depends on agonist binding in a reversible manner.

### Antagonists induce intermediate-size MR oligomers

Clinically relevant MR antagonists such as spironolactone, eplerenone or the recently approved non-steroidal antagonist finerenone induce nuclear translocation, although with slower kinetics (Amazit et al., 2015; Fejes-Toth et al., 1998; Gravez et al., 2013). We asked whether these antagonists also facilitate MR binding to HREs and what is the quaternary structure of the receptor under these conditions. Our results showed that all three antagonists induced MR binding to the MTTV array (Fig. 2). However, N&B revealed that MR oligomerization in the nucleoplasm and at the MMTV array does not reach the levels detected with agonists. Remarkably, there are clear differences between both antagonists. A saturating concentration of spironolactone produced an ε value of 2.04 in the nucleoplasm, consistent with a population mainly formed by MR dimers, and 4.43 at the MMTV array, consistent with tetramerization upon HRE binding (Fig. 2). Eplerenone showed lower ε, even at high antagonist concentration, with predominantly monomeric MR in the nucleoplasm (ε = 1.02-1.36, similar to the unstimulated receptor) and an ε of 2.31-2.75 at the MMTV array (Fig. 2). Finerenone also induced a predominantly monomeric MR in the nucleoplasm (ε = 0.90) and dimeric MR at the MMTV array (ε = 1.98) at saturating concentrations. These results suggest that the high-order oligomerization detected with saturating concentrations of agonists represent the fully active conformation of MR, while different antagonists produce intermediate steps in the building of the fully active oligomer.

**Figure 2.**
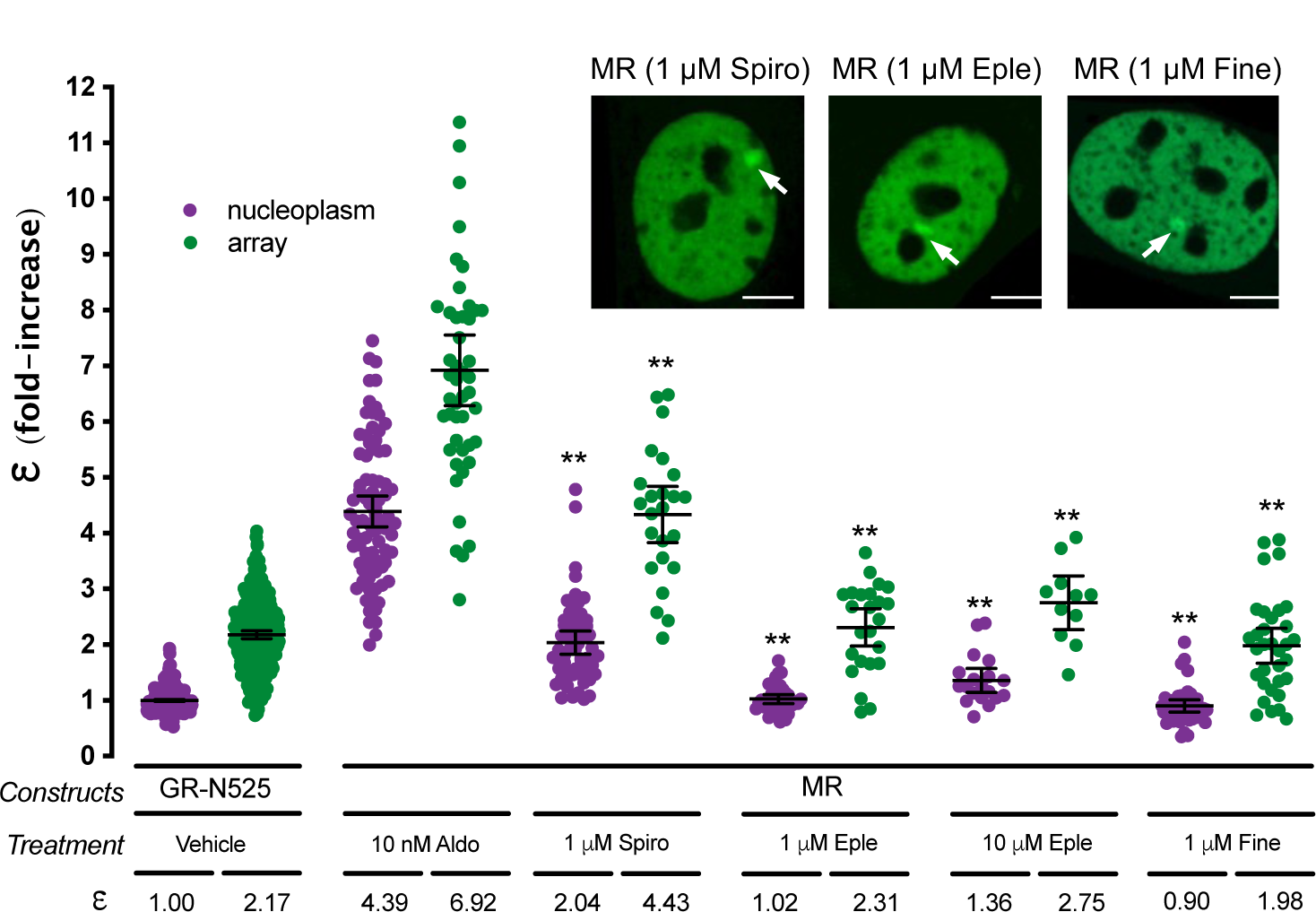
MR antagonists produce different MR quaternary configurations. Inset shows representative images of single cell nuclei expressing MR-GFP and treated with 1 µM spironolactone (Spiro), 1 µM eplerenone (Eple) or 1 µM finerenone. White arrows point to the MMTV array. Scale bars: 5 µm. Plot shows MR molecular brightness (ε) assessed using the N&B technique. To facilitate comparison, data from Fig.1 showing ε for GR-N525 and MR treated with 10 nM aldosterone are included. Data points correspond to ε obtained from a single nucleus (n = 490, 307, 82, 47, 55, 24, 37, 24, 19, 11, 41 and 33 cells in each condition, from left to right). Horizontal bars represent mean ± 95% CI. Each ε value was compared to its reference value (MR/10 nM Aldo in the nucleoplasm or at the MMTV array) using the Kruskal-Wallis test followed by Dunńs multiple comparisons correction. Symbols refer to statistical differences with MR/10 nM Aldo in the same compartment (nucleoplasm or MMTV array; **, p<0.01).

### MR and GR do not share the same dimerization interfaces

MR and GR evolved from a common corticoid receptor through gene duplication (Baker and Katsu, 2019; Bridgham et al., 2006) and have high sequence conservation in the DBD (94% identity) and moderate conservation in the LBD (57% identity; supplementary Fig. S2). Both domains harbor key determinants for GR oligomerization (Presman et al., 2016). The results described above uncover a profound difference between ligand-bound GR and MR quaternary conformation in the nucleus, probably reflecting different functionalities of the oligomerization interfaces between both receptors. To test this hypothesis, we analyzed the role of key residues in DBD and LBD in the process (Fig. 3A and 3B). To directly test the role of DNA binding, we first introduced the mutation C603S, which completely disrupts the first zinc finger in the DBD (Fig. 3A), eliminating the possibility of DNA binding (Cole et al., 2015; Pearce et al., 2002). This mutation produced a receptor that was unable to bind the MMTV array (Fig. 3C) and exists as a monomer in the nucleoplasm (Fig. 3D), suggesting that the effect of disrupting the P-loop is likely not limited to preventing DNA binding but has additional effects on the structure of the receptor.

**Figure 3.**
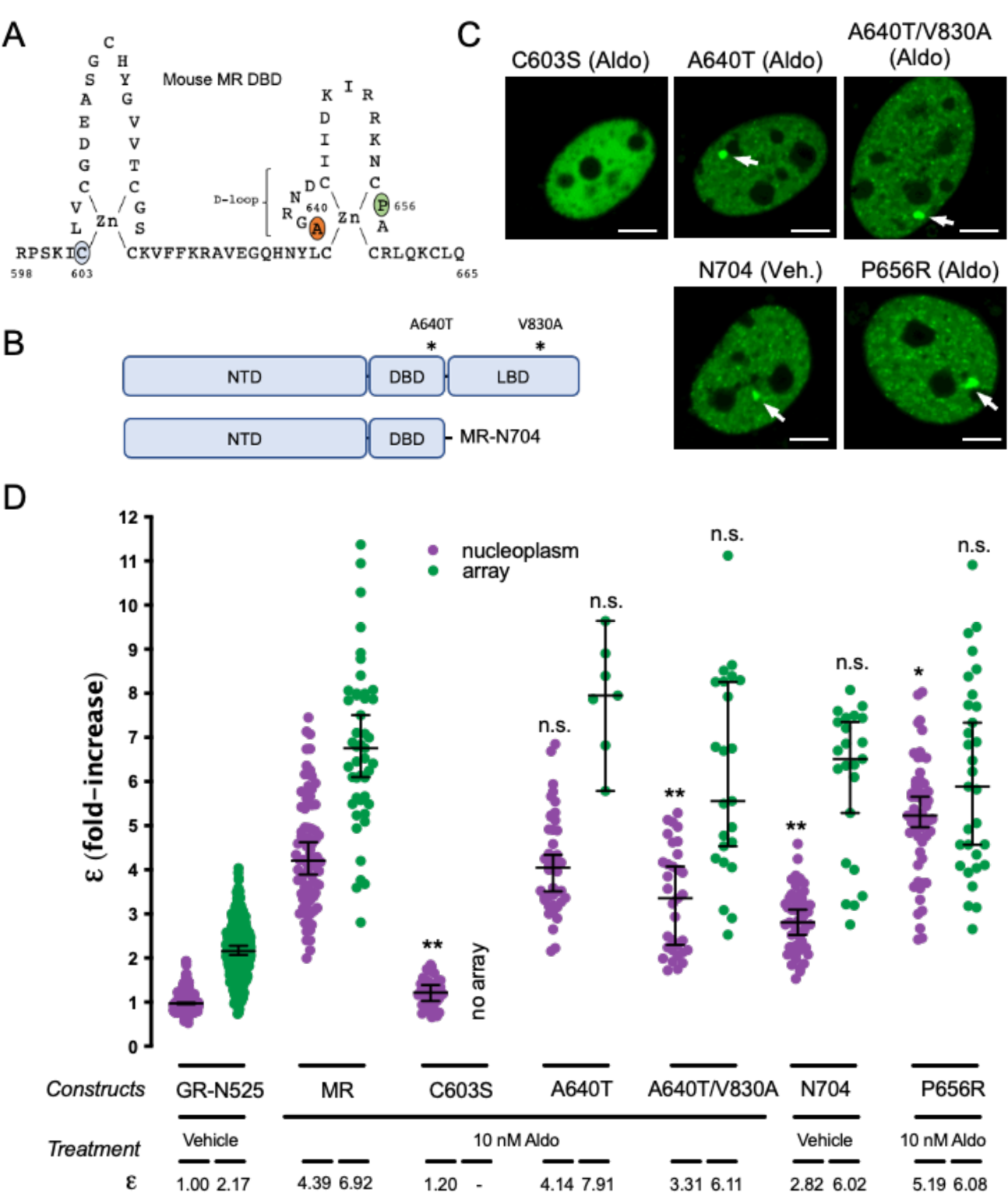
The role of predicted MR dimerization interfaces in quaternary structure formation. (A) Schematic representation of MR DBD. (B) Schematic representation of MR double mutant (A640T/V830A) and deletion generating construct MR-N704. NTD, N-terminal domain; DBD, DNA binding domain; LBD, ligand binding domain. (C) Representative images of single cell nuclei expressing the indicated constructs. White arrows point to the MMTV array. Scale bars: 5 µm. (D) MR molecular brightness (ε) assessed using the N&B technique. To facilitate comparison, data from Fig.1 showing ε for GR-N525 and MR treated with 10 nM aldosterone are included. Data points correspond to ε obtained from a single nucleus (n = 490, 307, 82, 47, 41, 44, 7, 33, 23, 51, 23, 57 and 31 cells in each condition, from left to right). To facilitate comparison, data from Fig.1 showing ε for GR-N525 and MR treated with 10 nM aldosterone are included. Horizontal bars represent mean ± 95% CI. Each ε value was compared to its reference value (wild type MR in the nucleoplasm or at the MMTV array) using the Kruskal-Wallis test followed by Dunńs multiple comparisons correction. Symbols refer to statistical differences with MR wild type in the same compartment (nucleoplasm or MMTV array; n.s., non-significant; *, p<0.05; **, p<0.01).

The DBD of GR contains a 5 amino acid sequence, the D-loop (supplementary Fig. S2) that has been proposed to be critical for dimerization (Dahlman-Wright et al., 1991). Mutation A465T in the D-loop of GR, commonly known as GRdim, was originally proposed to prevent DNA binding and dimerization of GR. Later work has shown that GRdim still forms dimers (Jewell et al., 2012; Presman et al., 2014), although its activity as transcriptional modulator is severely weakened (Johnson et al., 2021). Combination of A465T mutation with an additional mutation in the LBD (I634A) does produce a monomeric GR (GRmon; (Liu et al., 2020; Presman et al., 2016; Presman et al., 2014)). In the case of MR, the orthologous mutation equivalent to GRdim (A640T; Fig. 3A; supplementary Fig. S2) did not produce any effect on receptor oligomerization in the nucleoplasm or the MMTV array (Fig. 3D). The combination of point mutations A640T/V830A, equivalent to the double mutant A465T/I634A in GRmon, reduced values to 3.31 in the nucleoplasm but did not significantly affect oligomer size at the array (Fig. 3D). This result indicates that the function of these two well-described dimerization interfaces in GR is only partially conserved in MR, with a minor role in configuring its quaternary structure. To further test this idea, we deleted the whole LBD by truncating MR after amino acid 704 (MR-N704; Fig. 3B). MR-N704 is constitutively nuclear and binds the array in the absence of ligand (Fig. 3C), as described previously for the equivalent GR deletion, GR-N525 (Presman et al., 2016). MR-N704 produced reduced ε values in the nucleoplasm (2.82) but did not significantly affect oligomer size at the MMTV array (6) (Fig. 3D), similar to the A640T/V830A mutant. This stands in contrast to the equivalent deletion in GR, which produces monomers in the nucleoplasm and dimers at the MMTV array (Fig. 3; (Presman et al., 2016)), further indicating that the functionality of the classic GR dimerization interfaces is not fully conserved on MR.

To continue probing the role of the DBD and LBD dimerization interfaces, we introduced the mutation MR-P656R in the D-loop of the DBD (Fig. 3A), located in a residue that is fully conserved in GR (supplementary Fig. S2), where it mimics DNA binding, resulting in tetrameric receptors in the nucleoplasm with significantly enhanced binding to response elements inaccessible to wild type receptors (Paakinaho et al., 2019). N&B results showed that mutant MR-P654R shows a higher order oligomerization status in the nucleoplasm (ε = 5.19; Fig. 3D), approaching the value found at the array, although the effect is not as stark as the one found with GR (Presman et al., 2016). The activity of MR-P656R on enhancing the expression of well-known MR/GR target genes such as *Per1*, *Sgk1* or *Serpine1* was not significantly different from the WT construct (supplementary Fig. S3), similar to the effect of the equivalent mutation in GR, which does not significantly change the potency of the receptor but rather expands its set of target genes (Paakinaho et al., 2019).

### The NTD participates in MR tetramer formation in the nucleoplasm

Altogether, our analysis of highly conserved residues in the DBD and LBD of MR and GR that have been involved in GR dimerization indicates that their impact on MR is significantly lower, pointing to the involvement of other regions of the receptor in the formation of its quaternary structure. Given that the NTD shows low sequence conservation between MR and GR (<15% identity), we first deleted this entire domain, generating a truncated MR with the last 580 amino acids of the sequence (MR-580C, Fig. 4A). This construct showed an ε value of 1.63 in the nucleoplasm, indicating its essential role in ligand-induced, HRE-independent tetramerization of MR (Fig. 4B). In spite of this, MR-580C binds to the MMTV array (Fig. 4B), where it still forms higher order oligomers (ε = 6.61), almost indistinguishable from the wild type MR. This result suggests that tetramerization in the nucleoplasm is not a pre-requisite to form higher order oligomers at the MMTV array and also that MR DBD and LBD contain oligomerization interfaces that are triggered by HRE binding. Swapping the NTD of MR for the equivalent domain in GR (construct GR-N408/MR-580C, Fig. 4A) produced an intermediate oligomer size in the nucleoplasm (ε = 2.96, Fig. 4C), further confirming that ligand-induced tetramerization of MR requires both its NTD and LBD. This construct behaves almost normally at the MMTV array (ε = 6.38, Fig. 4C), again pointing to the importance of the DBD in higher order oligomerization. We also performed the opposite swap, creating a construct with the NTD of MR and DBD and LBD from GR (construct MR-N579/ GR-407C, Fig. 4A). Since aldosterone binds with lower affinity (Kd = 14 nM) and is a very poor activator of GR, we used 100 nM dexamethasone to stimulate this construct (Presman et al., 2016). The MR-N579/ GR-407C chimera produced an intermediate oligomer in the nucleoplasm (ε = 2.57), but still higher order quaternary organization at the MMTV array (ε = 7.89; Fig. 4C).

**Figure 4.**
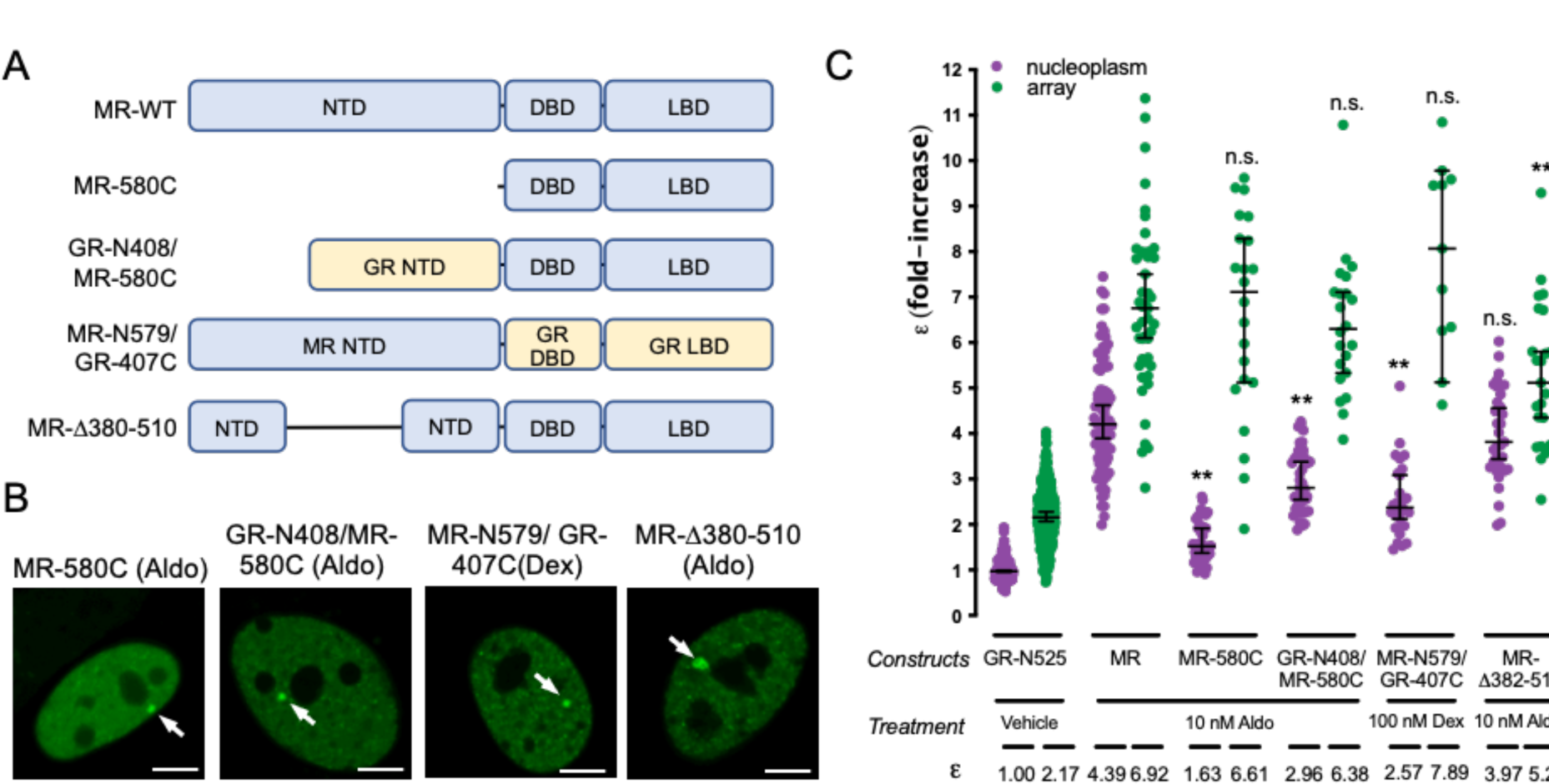
The NTD is essential for MR tetramerization in the nucleoplasm. (A) Schematic representation of MR deletion and domain swapping constructs. NTD, N-terminal domain; DBD, DNA binding domain; LBD, ligand binding domain. (B) Representative images of single cell nuclei expressing the indicated constructs. White arrows point to the MMTV array. Scale bars: 5 µm. (C) N&B results obtained with the indicated constructs. To facilitate comparison, data from Fig.1 showing ε for GR-N525 and MR treated with 10 nM aldosterone are included. Data points correspond to ε obtained from a single nucleus (n = 490, 307, 82, 47, 42, 22, 43, 22, 35 and 23 cells in each condition, from left to right). Horizontal bars represent mean ± 95% CI. Each ε value was compared to its reference value (wild type MR in the nucleoplasm or at the MMTV array) using the Kruskal-Wallis test followed by Dunńs multiple comparisons correction. Symbols refer to statistical differences with MR wild type in the same compartment (nucleoplasm or MMTV array; n.s., non-significant; **, p<0.01).

Taken together, our data suggest the possibility that MR has additional oligomerization determinants in the NTD. To further investigate this possibility, we generated a construct with deletion 382-510 (MR-Δ382-510 (Fig. 4A), which eliminates a region that has been previously implicated in a ligand-dependent N/C interaction in MR (Pippal et al., 2009), and should also disrupt the second part of the bi-partite activator function-1 domain (AF-1b; (Fuse et al., 2000; Tallec et al., 2003)). Remarkably, this deletion did not affect nucleoplasm tetramerization but clearly diminished oligomer size at the array (Fig. 4C), implicating this region in the formation of the final active conformation of MR.

### GR is able to incorporate into GR-MR heteromers, displacing MR subunits

GR and MR physically interact (Bigas et al., 2018; Jimenez-Canino et al., 2016; Liu et al., 1995; Nishi et al., 2004; Pooley et al., 2020; Rivers et al., 2019; Trapp et al., 1994), and their co-expression likely results in the modulation of both of their transcriptional programs (Bigas et al., 2018; Carceller-Zazo et al., 2023; Jimenez-Canino et al., 2016; Liu et al., 1995; Rivers et al., 2019; Trapp et al., 1994). The differences in quaternary organization between MR and GR raise an additional question. Would co-expressed MR and GR adopt a GR-like tetrameric conformation, an MR-like higher order oligomerization or a combination of both? To address this question, we performed N&B experiments in cells co-transfected with both receptors, eGFP-tagged MR with mCherry-tagged GR, recording data on the eGFP channel only. If GR incorporates into MR complexes, displacing some MR subunits, then MR’s ε value should drop in the presence of GR. On the contrary, if GR adds up to MR, then ε values should remain the same. Imaging of transfected cells showed that co-expression of both receptors produced simultaneous occupancy of the MMTV array (Fig. 5A). GR co-expression significantly reduced ε for MR, both in the nucleoplasm and in the array (Fig. 5B). Surprisingly, co-binding of both receptors to the MMTV array and the effect of GR lowering the apparent ε for MR occurred even when aldosterone was used as an agonist (Fig. 5A and 5B). It is worth noting that even though aldosterone does not activate GR at the concentration used in these experiments (10 nM), it does at least partially occupy the receptor (Kd = 14 nM; (Hellal-Levy et al., 1999)), as evidenced by its nuclear translocation and binding to the MMTV array even in the absence of MR (ε = 1.35 in the nucleoplasm and 2.15 at the MMTV array; supplementary Fig. S1B). Since these experiments use transiently transfected cells, the effect of co-expressed competing GR would be expected to vary depending on the relative proportion of MR and GR expressed in the cell.

**Figure 5.**
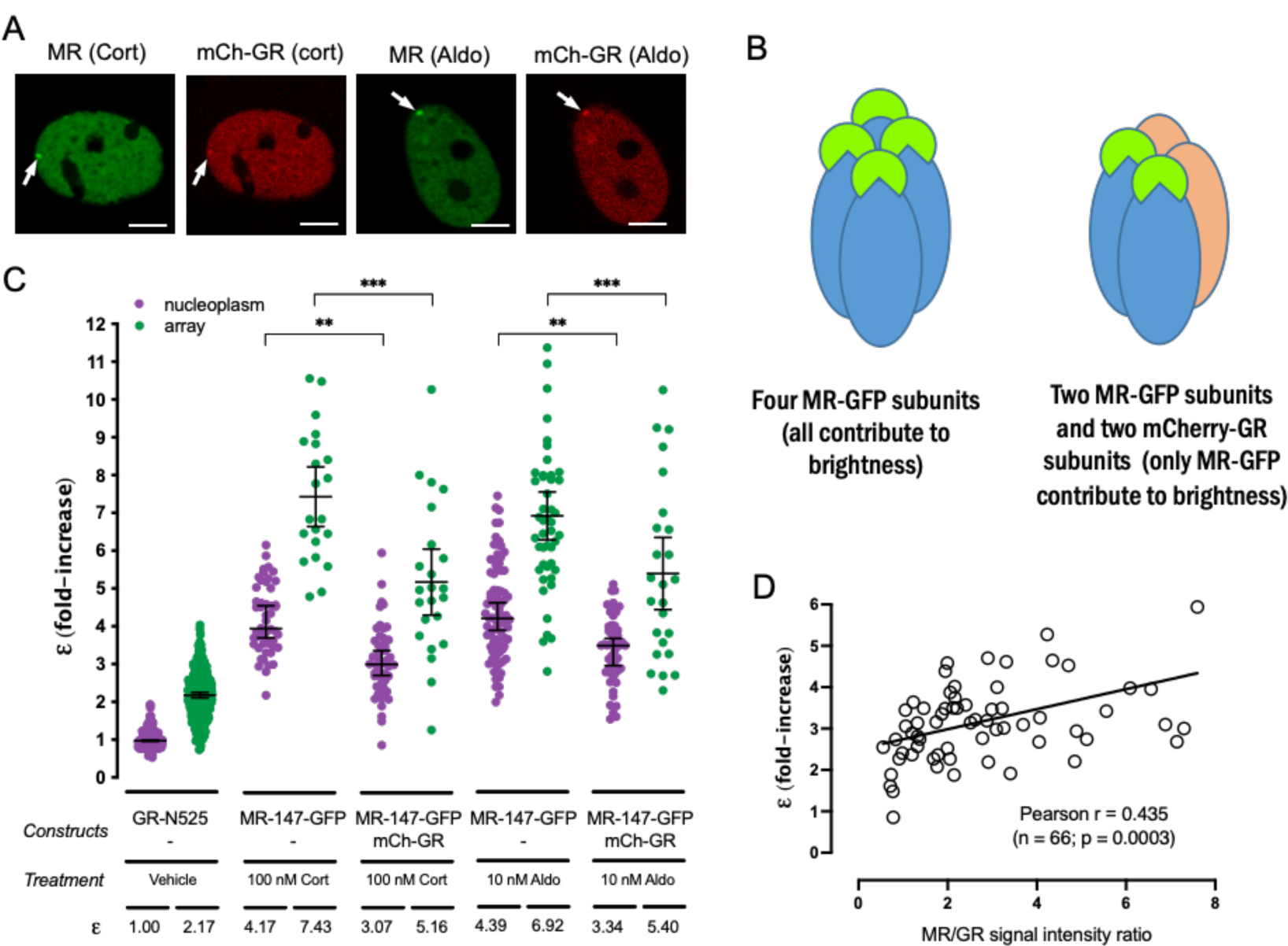
GR displaces MR subunits from the oligomer. (A) Representative images of single cell nuclei expressing the indicated constructs. mCh, mCherry. White arrows point to the MMTV array. Scale bars: 5 µm. (B) MR molecular brightness (ε) assessed using the N&B technique. To facilitate comparison, data from Fig. 1 showing ε for GR-N525 and MR treated with 100 nM corticosterone or 10 nM aldosterone are included. Data points correspond to ε obtained from a single nucleus (n = 490, 307, 50, 21, 50, 23, 82, 47, 50 and 25 cells in each condition, from left to right). Statistical analysis was performed using one-way ANOVA followed by Tukey test (results are shown only for two selected pairs; *, p < 0.05; **, p < 0.01; p < 0.001). (C) Linear correlation between eGFP-MR ε values and the ratio of MR/GR fluorescence intensity. Each dot represents an individual nucleus. Pearson lineal correlation coefficient was computed using Prism 9 (GraphPad).

Therefore, we measured the ratio of eGFP and mCherry intensities in each cell after each N&B recording and plotted it against ε, obtaining a positive correlation between both parameters (Fig. 5C). In conclusion, our data indicates that GR and MR can form heterocomplexes in live cells, wherein GR can displace some MR subunits, rendering a complex stoichiometry that requires further study.

## DISCUSSION

### Evolutionary conservation of quaternary structure and its impact on heteromerization

Here we used the N&B fluorescence imaging technique on living cells to study MR quaternary structure and its changes upon ligand binding and high-affinity interaction with HREs. Our results support a unique oligomeric conformation among the steroid receptor subfamily, with results compatible with tetramer formation induced by agonist binding, which generates larger oligomers after binding to specific sites in the DNA. This stands in contrast to the quaternary structure of GR (Presman et al., 2016), the closest relative of MR within the subfamily. It may be assumed that protein divergence during evolution is constrained by selective pressure to maintain protein structure, including stable protein-protein interactions forming quaternary arrangements (Echave and Wilke, 2017; Fornasari et al., 2007). It follows that proteins with closely related evolutionary origin and functions would be expected to share the same quaternary arrangement. In proteins that share very high sequence identity (> 90%), quaternary structure is almost always conserved (Marsh and Teichmann, 2015). However, proteins showing more moderate sequence identity tend to differ more in their quaternary structures. Indeed, it has been calculated that 30-40% identity correlates with a 70% probability of sharing a quaternary structure (Levy et al., 2008). For instance, the NSAR/OSBS subfamily of enzymes presents an overall sequence identity > 40%, but some of its members are dimers and some octamers (Odokonyero et al., 2014). It is also worth noting that the final quaternary structure of proteins might be underestimated due to bias in the techniques used to determine oligomeric states (Ali and Imperiali, 2005). Sequence conservation between MR and GR, and in general between steroid receptors, differs significantly between domains, with low conservation in the NTD (less than 15% identity), medium conservation in the LBD (approx. 60%) and highly conserved DBD (94% identity). Altogether it is not that surprising that MR and GR adopt different quaternary structures given that the lightly conserved NTD and the moderately conserved LBD play an important role in oligomer formation (Figs. 3 and 4).

It has long been known that nuclear receptors in the NR1 subfamily function by forming heteromers with RXR (De Bosscher et al., 2020). However, it is increasingly clear that NR3 subfamily receptors are also able to form so-called “atypical” heteromers, which include NR3 receptors interacting with NR1 receptors and RXR (De Bosscher et al., 2020). In addition, NR3 receptors form heteromers with members of the same subfamily. These include not only MR-GR interactions, but others such as GR association with PR, ER or AR (De Bosscher et al., 2020; Ogara et al., 2019), or MR interaction with ER (Barrett Mueller et al., 2014), just to name a few (De Bosscher et al., 2020). Taken together, this suggests that nuclear receptor crosstalk frequently involves formation of heteromers and that this is not hampered by the different quaternary conformations adopted by homomers, as confirmed by our data. This direct interaction model does not necessarily implicate direct binding of both kind of receptors to DNA, but may also involve tethering, looping or simply sequestration of one receptor by the other, preventing its genomic action. In the case of MR-GR interaction, it appears that they exert reciprocal effects on each other. It has recently been shown that MR modulates GR response to a synthetic glucocorticoid in mouse keratinocytes (Carceller-Zazo et al., 2023). In general, co-expression of MR and GR has been shown to alter glucocorticoid responses, although these effects appear to be cell-type and promoter-specific (Kiilerich et al., 2015; Liu et al., 1995; Mifsud and Reul, 2016; Ou et al., 2001; Rivers et al., 2019; Trapp et al., 1994). In addition, it is difficult to tease out whether transcriptional responses are primarily driven by GR, MR or both. It has also been proposed that MR may exert its effects not directly binding to DNA, but through tethering to GR (Rivers et al., 2019). There are few reports examining the role of GR on MR/Aldo function. Tsugita et al. used gene-reporter assays to show that GR co-expression is necessary for MR function, in a process likely involving receptor heteromerization and DNA binding (Tsugita et al., 2009), a model that we have recently confirmed investigating MR genome-wide binding and Aldo-regulated transcriptome (Johnson et al., 2023). It has recently been reported that abnormally high oligomeric forms of GR at the MMTV array, such as D647V that causes Chrousos disease, correlates with less transcriptional activity (Jimenez-Panizo et al., 2022). In this sense, it is tempting to speculate that MR’s higher oligomeric conformation is not optimal, and thus GR might increase MR activity (Johnson et al., 2023; Mifsud and Reul, 2016; Trapp et al., 1994; Tsugita et al., 2009) by reducing its stoichiometry. Nevertheless, whether the different reciprocal actions of MR and GR on their transcriptional activity reflect differences in the quaternary structure of the heterocomplexes requires further investigation.

### Mechanisms of oligomer formation

Results comparing binding of antagonists and agonists indicate that different ligands promote different average quaternary structures in the receptor population. Spironolactone, eplerenone and finerenone produced oligomers that are roughly multiples of two (Fig. 2). In fact, it is thought that an energetically-favorable, ordered pathway underlies formation of most protein complexes, although multiple parallel pathways are also possible (Marsh and Teichmann, 2015). Also, the subcomplexes formed during assembly of larger oligomers appear to correlate with evolutionary precursors (Marsh and Teichmann, 2014). It has been proposed that comparing homologous proteins with differing quaternary structures, such as the NR3 family of steroid receptors ((Presman et al., 2016) and the results presented here) can trace the evolution of a homomeric complex (Levy et al., 2008; Perica et al., 2012). Thus, it is tempting to speculate that building the final active conformation of MR involves an initial dimerization of soluble subunits upon ligand binding, possibly common to all steroid receptors, followed by stepwise doubling of oligomer size, which also appears to take place in other NR3 receptors, such as PR (Presman et al., 2016). The most potent agonist, aldosterone, does not generate an average oligomer size of 8, but reaches an average of 6.92 (Fig. 1), suggests that the population being observed could be an uneven combination of tetramers and octamers, with predominant presence of the latter.

Inter-subunit interfaces mediating the different steps in forming MR tetramers or higher order oligomers remain to be studied in detail. Our data indicate that all three domains (NTD, DBD and LBD) play a role, with complex interactions between them. In general, it appears that each domain has a modest contribution on its own, as reflected by small decreases in ε when they are disrupted by deletion of point mutations in known GR dimerization interfaces. The only exception is mutation C603S, which completely disrupts the first zinc-finger domain in the DBD. This mutation renders MR monomeric and unable to bind DNA (Cole et al., 2015; Pearce et al., 2002). However, it is not possible to infer that the DBD is the essential domain for DNA oligomerization, since the impact of this mutation on the structure of the receptor may be wider and also, it precludes DNA binding and possibly allosteric interactions between the DBD and LBD, proposed to play an important role in GR tetramerization (Presman et al., 2016; Presman and Hager, 2017).

### Functional impact of oligomerization

The correlation between agonist and antagonist binding and the size of the MR oligomer strongly suggests that higher order oligomerization detected only after agonist and DNA binding represents an active conformation of the receptor. On the other hand, and as discussed above, the possibility remains that GR modulation of MR stoichiometry may increase the activity of the latter. In the case of GR, the “tetra” mutation, P481R, stabilizes the tetrameric conformation of the receptor (Presman et al., 2016), allowing functional dissection of tetramerization, which has been shown to potently drive chromatin interactions and transcriptional activity (Paakinaho et al., 2019). The equivalent mutation in MR, P656R, does increase oligomer size in the nucleoplasm from 4.39 to 5.19, but this is a much more modest change that does not appear to change transcriptional regulation of endogenous genes (supplementary Fig. S3). It remains to be examined whether this small increase would produce a genome-wide effect changing MR chromatin binding and/or transcriptional outcomes. Ideally, further investigating the mechanisms of MR oligomer formation will provide tools to manipulate oligomer size in the presence of agonists, making it easier to assess its functional importance.

### Conclusions

In summary, we have shown that MR adopts a distinct quaternary structure, further supporting a model where each steroid receptor adopts different conformations that are not determined by their evolutionary relationships. These differences do not appear to preclude formation of heteromeric complexes between NR3C receptors. Large MR oligomers at the MMTV array are reached only with agonists, strongly suggesting a relationship between oligomer size and the final active conformation of the receptor. Structural determinants of MR oligomerization appear complex, with participation of all three main domains. These results have important implications for the pharmacological modulation of steroid receptor signaling.

## MATERIAL AND METHODS

### Plasmids constructs and mutagenesis

A fully functional mouse MR fluorescent derivative with insertion of enhanced green fluorescent protein (eGFP) after amino acid 147 has been previously described (Aguilar-Sanchez et al., 2012). N-terminus eGFP tagged wild type mouse GR or truncated mutant GR-N525, lacking the entire LBD, (eGFP-GR and eGFP-GRN525, respectively) have been previously described (Meijsing et al., 2007; Presman et al., 2016). A plasmid expressing mouse GR tagged in the N-terminus with mCherry was developed by amplifying mouse GR coding sequence and in-frame cloning in pmCherry-C3 (Clontech). Point mutations and deletions were introduced using the Quickchange XL mutagenesis kit (Agilent) following the manufactureŕs instructions. Domain swapped mouse MR/GR constructs were constructed by amplification of the relevant fragments from donor plasmids and directional cloning using ligation-free In-Fusion technology (Clontech) in PCR-mediated linearized vectors. PCRs were performed using high-fidelity Pfu ultra II polymerase (Agilent). All constructs and mutations were confirmed by DNA sequencing.

### Cell culture, transfection and treatment with ligands

The cell lines used in this study originally derive from mouse mammary carcinoma cell line C127 (RRID: CVCL_6550), which were originally modified introducing approximately 200 copies of a tandem array of the Harvey viral ras (MMTV-v-Ha-ras) reporter, which contains several HREs (McNally et al., 2000). This cell line was then modified using CRISPR/Cas9 to knockout the endogenous expression of GR, as previously described (Paakinaho et al., 2019). When indicated, a cell line with stable integration of a plasmid containing eGFP-MR driven by the CMV promoter was used. This cell line was developed using the CRISPR/Cas9 procedure, as previously described (Paakinaho et al., 2019). Cells with plasmid insertion were selected by puromycin treatment followed by fluorescence-activated cell sorting (FACS). Since expression of MR in the sorted polyclonal cell population declined over time, we further selected stable lines by single-cell cloning. eGFP-MR expression was confirmed by confocal microscopy previously described (Jimenez-Canino et al., 2016). All cell lines were grown in Dulbecco’s modified Eagle’s medium (DMEM), 10% fetal bovine serum (FBS), sodium pyruvate, nonessential amino acids and 2 mM glutamine. Culture medium and reagents were obtained from Gibco, except FBS, which was from Gemini. Culture medium also contained 5 μg/mL tetracycline (Sigma–Aldrich) to prevent expression of a stably integrated eGFP-GR (Presman et al., 2014). Cells were maintained at 37C and 5% CO2 in a humified incubator. Forty-eight hours before experiments, cells were plated in 2-well borosilicate glass chambers (Nunc™ Lab-Tek™ II, ThermoFisher #155379) in DMEM supplemented with 10% charcoal/dextran-treated FBS (CS-FBS). Next day, cells were transfected using Jetprime (Polyplus) according to the manufactureŕs instructions. Ligands were obtained from Sigma-Aldrich (aldosterone, corticosterone, spironolactone and eplerenone) or MedChemExpress (finerenone). On the day of the experiment, cells were treated for 1h with ligands dissolved in ethanol (vehicle) and added to the medium at the final concentration indicated in each experiment. Ligand concentrations were generally chosen to fully saturate the receptor (Amazit et al., 2015; Hellal-Levy et al., 1999), except when indicated. Hormone washout was performed by three consecutive 10-minute washes with prewarmed hormone-free complete medium with CS-FBS, followed by incubation in the same medium for 4h. All ligands were obtained from Sigma-Aldrich.

### Number and brightness (N&B) analysis

N&B was performed as previously described (Presman et al., 2016; Presman et al., 2014). Briefly, cells were placed in an environmentally controlled chamber in a Zeiss LSM780 (CCR Confocal Microscopy Core Facility, NIH, Bethesda, Maryland, USA) or LSM980 (Hospital Universitario NS Candelaria, Tenerife, Spain) confocal microscopes and maintained at 37°C and 5% CO_2_ throughout the duration of the experiment. After approximately 30 min of equilibration, single nuclei were imaged using a 63X oil immersion objective (N.A. = 1.4). Cells were imaged between 30 min and 2h after addition of each ligand. Fluorescence excitation was performed with a multi-line argon laser tuned at 488 nm and detection was performed with a gallium arsenide phosphide detector set in photon-counting mode. For each nucleus, stacks of 150 images (256 x 256 pixels) from a single plane were taken, using a pixel size of 80 nm and a pixel dwell time of 6.3 µs. In the case of recordings performed with the LSM980 microscope, we collected stacks of 120 images with a pixel dwell time of 8.19 µs. Recording conditions ensured sampling of independent populations of molecules (Mikuni et al., 2007; Presman et al., 2016; Presman et al., 2014). Data analysis was performed using Globals for Images · SimFCS 2 software (Laboratory for Fluorescence Dynamics, University of California, Irvine, CA). Pixels were classified as nucleoplasm or MMTV array according to their intensity values. Quality control for analysis followed our previously described guidelines (Presman et al., 2016; Presman et al., 2014). Average fluorescence intensity () and variance (σ2) were calculated for each pixel along the image stack. The ratio of σ2 to provides the apparent brightness (B). Real brightness (ε) was calculated as B – 1 (Presman et al., 2016; Presman et al., 2014). Each experimental condition was repeated independently two to five times. Results from all experiments were pooled after normalizing with the internal monomeric control (GR-N525). When indicated, mCherry-GR was co-transfected with eGFP-MR and the average intensity of mCherry fluorescence was recorded for the nucleus of interest before performing the N&B experiment.

### RNA Isolation, qPCR and RNA-seq analysis

Cells cultured for at least 24h on CS-FBS supplemented culture medium were treated with vehicle, 10 nM aldosterone or 100 nM corticosterone for 2 h. Total RNA was purified using a commercially available kit (Macherey-Nagel NucleoSpin RNA isolation) that includes an in-column DNase digestion step. Purified RNA was quantified using spectrophotometry and frozen at −80°C. Single-stranded cDNA was synthesized from 1 µg of total RNA as template using a commercially available kit (iScript cDNA Synthesis Kit, Biorad). RT-qPCR analysis of nascent mRNA abundance was performed in duplicate using iQ SYBR Green Supermix (Biorad #1708880) in a Biorad CFX96 machine. Primers for the amplification of nascent Per1, Sgk1 and Serpine1 were as described (Johnson et al., 2021).

### Statistical analysis

Statistical analysis was performed using Prism 9 (GraphPad). Gaussian distribution of N&B ε values was performed using the D’Agostino-Pearson omnibus K2 normality test. When all conditions analyzed passed the normality test, parametric tests were used (t test when comparing only a selected pair of conditions; one way ANOVA followed by Tukey test when comparing more than two conditions). When not all compared conditions passed the normality test, a non-parametric test was used (Kruskal-Wallis followed by Dunńs multiple comparisons correction).

## Acknowledgments

This research was supported by the Intramural Research Program of the NIH, National Cancer Institute, Center for Cancer Research. DAdlR was supported by grants PID2019-105339RB-I00 (funded by MCIN/AEI/10.13039/501100011033 and “ERDF A way of making Europe”), and PRX18/00498 (funded by *Programa Estatal de Promoción del Talento y su Empleabilidad en I+D+i, Subprograma Estatal de Movilidad, del Plan Estatal de I+D+I,* MICINN, Spain). BA-P was supported by pre-doctoral fellowship BES-2017-082939 (funded by MCIN/AEI/ 10.13039/501100011033 and by “ESF Investing in your future”). D.M.P. was supported by CONICET and the Agencia Nacional de Programación Científica y Tecnológica [PICT 2019-0397 and PICT 2018-0573]. The authors declare that they do not have competing interests.

## Author contribution

G.F.: conceptualization, methodology, investigation, validation, formal analysis.

T.A.J.: conceptualization, methodology, validation, formal analysis, investigation, writing, visualization. B.A.-P.: investigation.

D.M.P.: methodology, validation, writing.

G.L.H.: conceptualization, formal analysis, writing, supervision, funding acquisition.

D.A.d.l.R.: conceptualization, investigation, formal analysis, writing, visualization, supervision, funding acquisition.

## Data and materials availability

All data and materials are available upon reasonable request.

