## Supplementary Figures S1-S3 for "The mineralocorticoid receptor forms higher order oligomers upon DNA binding"

**SUPPLEMENTARY FIGURES, Fettsweis et al.**

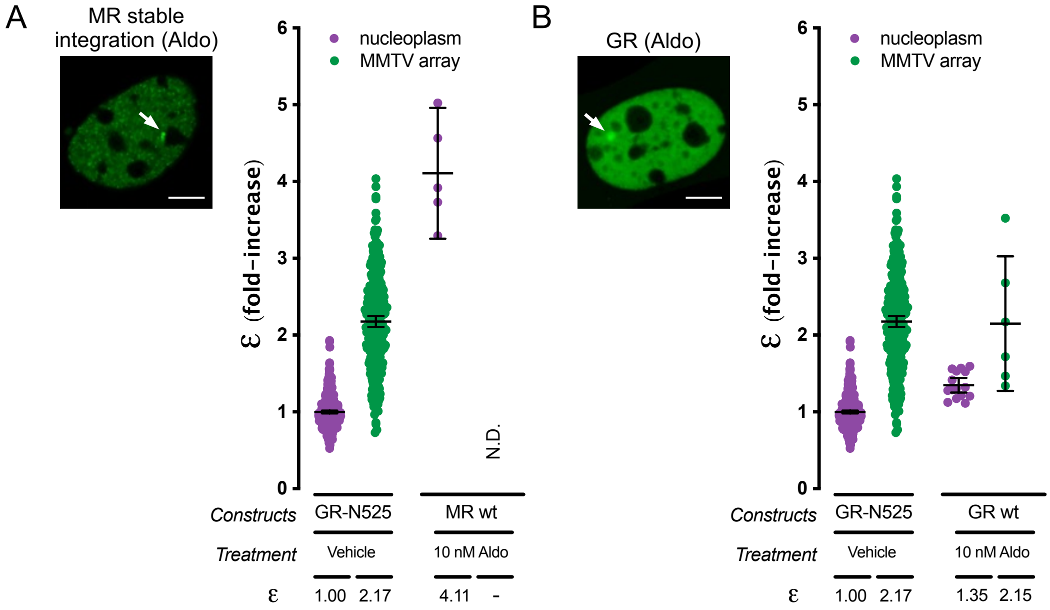


**Supplementary Figure S1**. (A) Representative image and molecular brightness (ε) in the nucleoplasm of cells stably expressing eGFP-tagged mouse MR (MR). We were unable to record from the MMTV array due to low levels of receptor expression. Individual dots represent values from one cell (n = 490, 307, 5; N.D., not determined). (B) Representative image and molecular brightness (ε) obtained from cells expressing eGFP-tagged wild type mouse GR (GR) and treated with 10 nM aldosterone for 1h (n = 490, 307, 15, 6). White arrows point to the MMTV array. Scale bars: 5 $\mu$m. To facilitate comparison, data from Fig.1 showing ε for GR-N525 in the nucleoplasm and MMTV array are shown.

A

```
GR DBD   423 PPPKLCLVCSDEASVCHYGVLTCGSCKVFFKRAVEGQHNYLCAGRNDCIIDKIRRKNCPACRYRKCLQ 490
MR DBD   598 RPSKICLVCGDEASGCHYGVVTCGSCKVFFKRAVEGQHNYLCAGRNDCIIDKIRRKNCPACRLQKCLQ 665
              * *:****.**** *****:***************************************** :****
```

B

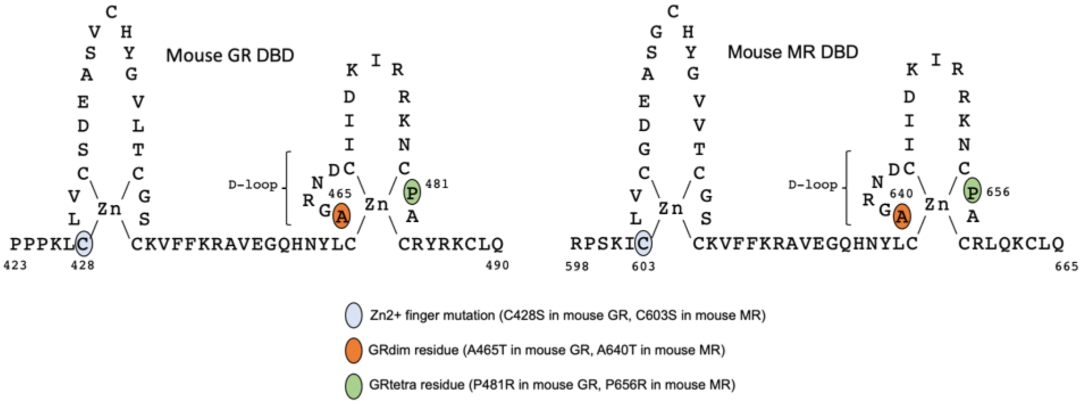


C

```
GR LBD   526 -----------------IVPAALPQLTPTLVSLLEVIEPEVLYAGYDSSVPDSAWRIMTTLNMLGGRQ 576
MR LBD   705 TYIAPTKEPSVNSALVPQLASITRALTPSPSMILENIEPEIVYAGYDNSKPDTAESLLSTLNRLAGKQ 772
                           : :    ***:   :** ****::*****.* **:*  :::*    ** *.*:*

GR LBD       VIAAVKWAKAIPGFRNLHLDDQMTLLQYSWMFLMAFALGWRSYRQASGNLLCFAPDLIINEQRMTLPC 644
MR LBD       MIQVVKWAKVLPGFKNLPLEDQITLIQYSWMCLSSFALSWRSYKHTNSQFLYFAPDLVFNEEKMHQSA 840
             :* .*****.:***:** *:**:**:***** * :***.****::..::* *****::**::*   .

GR LBD       MYDQCKHMLFISTELQRLQVSYEEYLCMKTLLLLSSVPKEGLKSQELFDEIRMTYIKELGKAIVKREG 712
MR LBD       MYELCQGMRQISLQFVRLQLTFEEYSIMKVLLLLSTVPKDGLKSQAAFEEMRTNYIKELRKMVTKCPN  908
             **: *: *  ** :: ***:::***  **.*****:***:*****  *:*:* .***** * :.*  .

GR LBD       NSSQNWQRFYQLTKLLDSMHDVVENLLSYCFQTFLDK-SMSIEFPEMLAEIITNQIPKYSNGNIKKLL 779
MR LBD       SSGQSWQRFYQLTKLLDSMHDLVNDLLEFCFYTFRESQALKVEFPAMLVEIISDQLPKVESGNAKPLY 976
             .*.*.****************:*::**.:** ** :. ::.:*** **.***::*:** ..** * *

GR LBD       FHQK 783
MR LBD       FHRK 980
             **:*
```

**Supplementary Figure S2**. (A) Sequence comparison between mouse GR and MR DBDs. Highlighted residues were mutated during this study. (B) Schematic representation of mouse GR and MR DBDs, indicating key residues mutated in MR during this work. (C) Sequence comparison between mouse GR and MR LBDs. Residues GR-I634, part of an LBD-LBD dimerization interface (8) and conservative change MR-V830 are highlighted. Alignments were performed using Clustal Omega (71) and sequences with GenBank accesion numbers AAI29913 and AAI33714 for GR and MR, respectively.

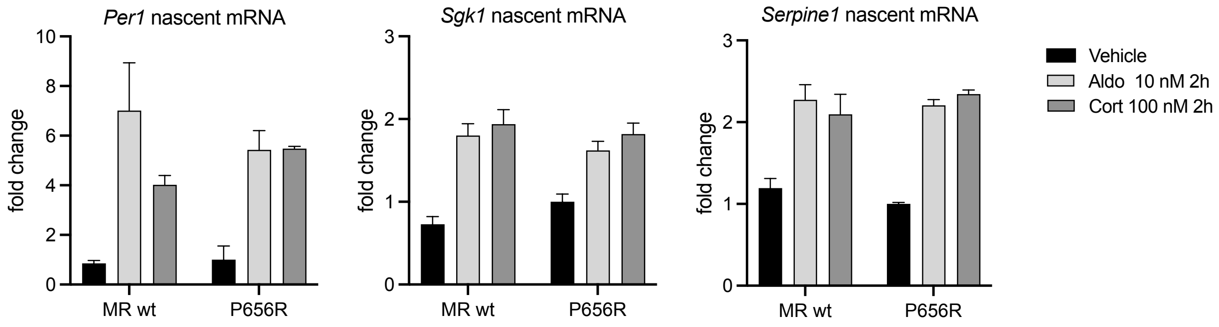


**Supplementary Figure S3**. RT-qPCR performed on three MR up-regulated genes in cells expressing wild type MR or MR mutant P656R. Cells were treated with vehicle, with 10 nM aldosterone or 100 nM corticosterone for 2h. Plots show fold changes in the indicated nascent mRNA abundance compared to MR-P656R treated with vehicle (n = 2).
